# Interpretable Visualization of Scientific Hypotheses in Literature-based Discovery

**DOI:** 10.1101/2021.10.29.466471

**Authors:** Ilya Tyagin, Ilya Safro

## Abstract

In this paper we present an approach for interpretable visualization of scientific hypotheses that is based on the idea of semantic concept interconnectivity, network-based and topic modeling methods. Our visualization approach has numerous adjustable parameters which provides the domain experts with additional flexibility in their decision making process. We also make use of the Unified Medical Language System metadata by integrating it directly into the resulting topics, and adding the variability into hypotheses resolution. To demonstrate the proposed approach in action, we deployed end-to-end hypothesis generation pipeline AGATHA, which was evaluated by BioCreative VII experts with COVID-19-related queries.

## 1 INTRODUCTION

Staying current in the fast growing amount of publicly available scientific literature is one of the main challenges the researchers are struggling with. Various machine learning models and artificial intelligence systems try to automatically connect previously disconnected concepts and formulate novel scientific hypotheses to accelerate the research pace by minimizing the time researchers spend on deriving information chains or more complicated patterns. However, interpretability of the results obtained by the literature based discovery and hypothesis generation systems has not yet reached its advanced stage representing in fact a very limited number of techniques.

Interpretation and evaluation of the literature based discovery and hypothesis generation systems is extremely difficult because the correct results (i.e., novel scientific discoveries or insights) are unknown if the system is used for proactive research. Hypothesis generation should not be confused with the information retrieval because the primary goal of the former is discovering novelty while the latter are looking for existing information. In the same time, retroactive verification is also not easy due to the challenges of perfect separation of test and training data. Some systems are designed with the interpretability in mind in the first place (e.g., MOLIERE [1]), but the large-scale experiments show that they are outperformed by less interpretable modern deep learning-based counterparts [2] even if designed using similar principles. To fill the gap between interpretable and state-of-the-art performing hypothesis generation systems, we propose a method to visualize hypotheses generated by the knowledge graph based systems with the use case demonstrated on AGATHA, the graph mining and transformer based model.

The proposed visualization is based on high interconnectivity of the biomedical concepts, i.e., typically, there is a very high chance that a pair of biomedical concepts is meaningfully connected in semantic knowledge networks. In these graphs, nodes and edges represent biomedical concepts (e.g., terms, genes, diseases, and papers) and semantic connections between them (e.g., co-occurrence in the paper, and experimental evidence of relation extracted from some database), respectively. To represent the nature of this data we use a network-based approach: all concepts are put in a large knowledge network in which the problem of finding a connection between any pair of them is formulated as finding a shortest path in a graph. Because such paths are usually short and not too informative, it is then *enriched with a cloud of textual information* located around it in this network, such that the connection between source and target terms is represented by a corpus of documents. Depending on the implementation and model, these documents might be represented by sentences or the entire abstracts. After the relevant documents are collected, *the topic modeling* is applied which results in a fuzzy clustering of the documents by topics. Each topic is, in turn, represented by documents and keywords. This fuzzy clustered representation of information is our explainability tool. The resulting topic model would represent a summary of documents related to both input terms, which can significantly improve the understanding the nature of their potential connection as it is expected to be not trivial and cannot be obtained with information retrieval systems.

## 2 BACKGROUND AND RELATED WORK

The main baseline and point of comparison for the proposed visualization approach is MOLIERE system released in 2017 [1]. It is a general purpose hypothesis generation system, which, at it was mentioned, was made with the interpretability in the first place. Its main concept of using shortest paths may look obsolete for highthroughput queries ran *on scale* and it is undoubtedly inferior to deep learning methods in terms of query execution speed, but this approach still holds when it comes to the visualization due to its natural explicitness and conceptual clarity.

In the proposed approach shortest path concept is used *for visualization purposes only*. It helps us to obtain high quality ranking keeping the running times reasonable and perform time-consuming visualization step only when it is truly necessary. In addition, the main unit of information is changed from a full abstract to a single sentence, significantly increasing the knowledge base size and the overall problem complexity.

Another important point of comparison is that MOLIERE resulting topics are static once generated, whereas in AGATHA Semantic Visualizer we perform generating topics *on the fly*. Topic modeling occurs during the browser session and only takes a couple of seconds to run. It is a significant improvement because the result may vary drastically with changing the underlying parameters.

## 3 METHODS AND IMPLEMENTATION

The entire visualization pipeline consists of three major parts: 1) AGATHA semantic graph construction, 2) visualization preparation and 3) interactive session. We describe each of these parts in this section.

### 3.1 AGATHA Semantic Graph Construction

We start with building a large heterogeneous graph, which is used as a data source for both inference and visualization purposes. For that we perform text pre-processing stage, which includes splitting all MEDLINE abstracts into sentences and extracting different semantic structures from these sentences. We note that according to our experiments processing the full text papers requires significantly more computational and time resources which does not necessarily improves the prediction quality [3]. The graph contains nodes that correspond to the following types of semantic objects:

- Sentences (*s* type)
- Lemmas (*l* type)
- N-grams (*n* type)
- Entities (*e* type)
- Predicates (subject-verb-object triples) (*p* type)
- MeSH and UMLS terms (*m* type)

The edges between nodes are added based on the following rules:

- Semantic structure (e.g. UMLS term or n-gram) is present in the sentence.
- Sentences have similar embedding vector representations.
- Sentences are from the same abstract and follow each other.
- Predicates are connected to both subject and object terms they contain.

As a result, we have a network containing hundreds of millions of nodes and billions of edges. The diagram of the network illustrated with a sample sentence and its extracted semantic objects is shown on Figure 1. For more information we refer to the full description of AGATHA model [2, 4].

**Figure 1:**
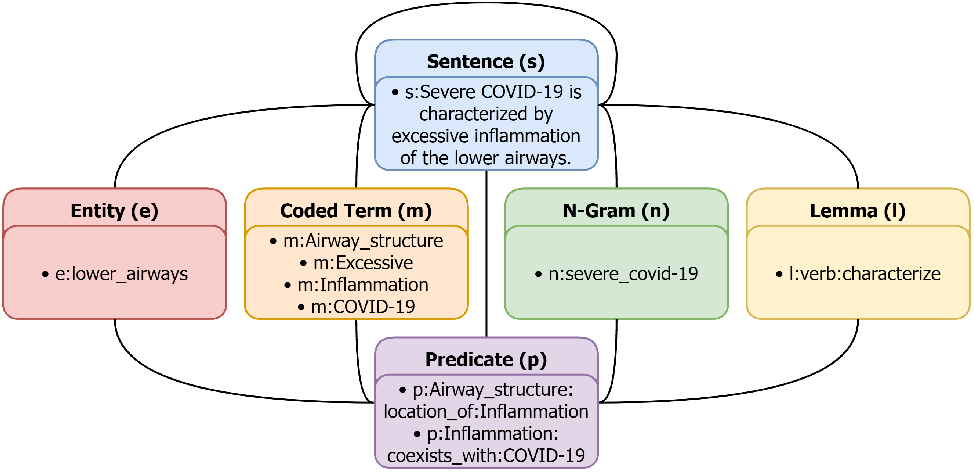
AGATHA Semantic Graph Structure.

To obtain a low-dimensional representation of this network, we perform *graph embedding* stage with PyTorch BigGraph Graph Embedding System [5], which puts every node in a unified vector space of dimensionality *d* = 512, such that similar nodes are clustered together.

After the graph is constructed and embedded, we train AGATHA deep learning model, which main purpose is to predict plausible term-term connections and for each input query generate a pairwise confidence score in the unit interval. The input format for the system is a pair of terms (source and target), which both should be represented as UMLS CUIs (Concept Unique Identifiers).

### 3.2 Visualization Preprocessing

In order to generate a visualization relevant to the input query, the first step is to obtain the subset of nodes in the AGATHA semantic graph, which would be *relevant to both source and target terms*. For that we first locate source and target terms in the semantic graph and find a shortest path between them. From our experience, these paths are typically of length of 3-5. We should admit that this shortest path is not representative, it does not contain enough information about the query connection and is not very useful by itself. However, the information around or within a certain proximity of the path nodes can be used to represent the subset of data relevant to both source and target terms.

Specifically, nodes of type *s* (sentence) are collected in the first place because they are considered as a corpus of documents for topic modeling and they are further processed during the interactive session. Another piece of information collected from the graph is sentence neighbors or semantic structures contained in the extracted sentences. These structures are used to form a vocabulary during the visualization part. We also keep track of the embeddings of these terms and add them to the visualization input.

### 3.3 Interactive Session

When all the necessary data is collected (that is, source and target terms, AGATHA confidence score, input corpus and its adjacent nodes with their vector representations), interactive session can be started. Its main idea is to *summarize the provided corpus and represent it in a network format*. For that two main components are used: Latent Dirichlet Allocation (LDA) and k-Nearest Neighbors (KNN).

The **LDA** [6] model is a statistical approach to find a set of implicit (unobserved) fuzzy clusters in the input dataset. In case of natural language processing, LDA is a way to represent the input corpus as a user-parametrized number of topics that cluster the documents in an explainable way. Each document is represented as a mixture of topics and each topic is represented as a mixture of keywords used in the text corpus. After applying LDA to the input corpus, we obtain a list of topics, which we then feed into the KNN algorithm.

An important addition to the LDA algorithm that we propose is a *bias towards terms of preferred category*. It was mentioned earlier that the vocabulary consists of semantic structures discovered in sentences, not ordinary tokens. It significantly improves the vocabulary quality and makes topics more sensible overall. This strategy also makes possible to track and tune *what vocabulary terms LDA should prioritize*.

A document *d* in LDA model is represented as a bag of vocabulary terms *w*_*i*_, where *i* - number of times a term *w* occurs in *d*. This approach treats all terms equally based on their raw document occurrences, but it can be trivially extended if additional information about terms *w* is available. In our case, *w* may belong to the following categories:

- Lemmas (l type)
- N-grams (n type)
- Entities (e type)
- MeSH and UMLS terms (m type)

These categories are presented in the order of increasing specificity: lemmas and n-grams are generally more broad than entities and MeSH/UMLS terms. Moreover, MeSH and UMLS terms come with semantic types as part of UMLS metathesaurus ^1^.

Given all that, we can now adjust the bag-of-words document representation to prioritize the preferred categories of vocabulary terms: now each document *d* is represented as a bag of vocabulary terms 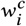, where *c* ∈ {*l, n, e, m*} - term category. When *c* = *m*, it is possible to identify a semantic type of a specific vocabulary keyword.

If one would like to prioritize a specific keyword category (for example, n-grams), we simply duplicate the keywords belonging to this category in bag-of-words representation as many times as it may be needed to bias LDA model towards uncovering topics containing the desired keyword categories.

Semantic types make this procedure even more useful: they allow a domain expert to pick a specific set of keyword types of interest (e.g. genes or diseases) and force LDA to generate results with higher likelihood of these terms appearing in the end topics.

The **KNN** stage takes place when the result topical network is constructed. KNN algorithm takes a set of vectors as input and produces a network of closest *k* neighbors for these vectors. In order to obtain the vectors for each topic, we represent them as centroids of keywords they contain:

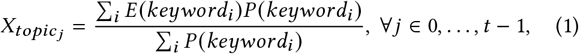

where 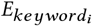 is an embedding for *i*-th keyword and 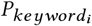 is a probability of *i*-th keyword being in *topic*_*j*_, *t* - number of topics in LDA model. In addition, 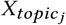 is a vector from the same space as the vocabulary terms, thus it can be compared to them. It is important because the topical KNN network would contain *t* + 2 nodes: *t* topic centroids as well as source and target terms. The main goal of building the topical network is to be able to connect source and target terms through intermediate topics using the shortest path strategy. A shortest path would represent a *hypothesis*: a suggested set of topics appearing “between” provided terms of interest. Topical graph techniques are discussed in details in [7].

To display results in interface, UMAP [8] algorithm is used. It reduces dimensionality for both centroid and term vectors from *d* = 512 to *d*_*red*_ = 2, which allows to draw them on screen. UMAP is chosen because it is known to preserve vectors topological features, such as global structure. UMAP-obtained vectors are used for *visualization-only purposes*: topic centroids and KNN network are constructed using the original *d* = 512 vectors obtained with PTBG system.

## 4 RESULTS

This particular work focuses on the visualization part of the hypothesis generation process, thus the results do not include any numerical validation. The AGATHA system which this work is based on was thoroughly evaluated via time-slicing strategy demonstrating state-of-the-art results in different biomedical domains. For more details please refer to the original papers [2, 4].

In this section we would like to demonstrate visualization features and summarize the feedback obtained during the BioCreative VII IAT evaluation process.

### 4.1 Deployment Details

To improve the overall system user experience, we deployed an AGATHA end-to-end pipeline, which is made of multiple building blocks and contains several important components. It is summarized on Figure 2.

**Figure 2:**
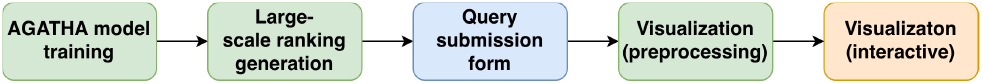
AGATHA end-to-end pipeline. Colors: green: computational server; blue: Google servers; orange: visualization server.

**Figure 3:**
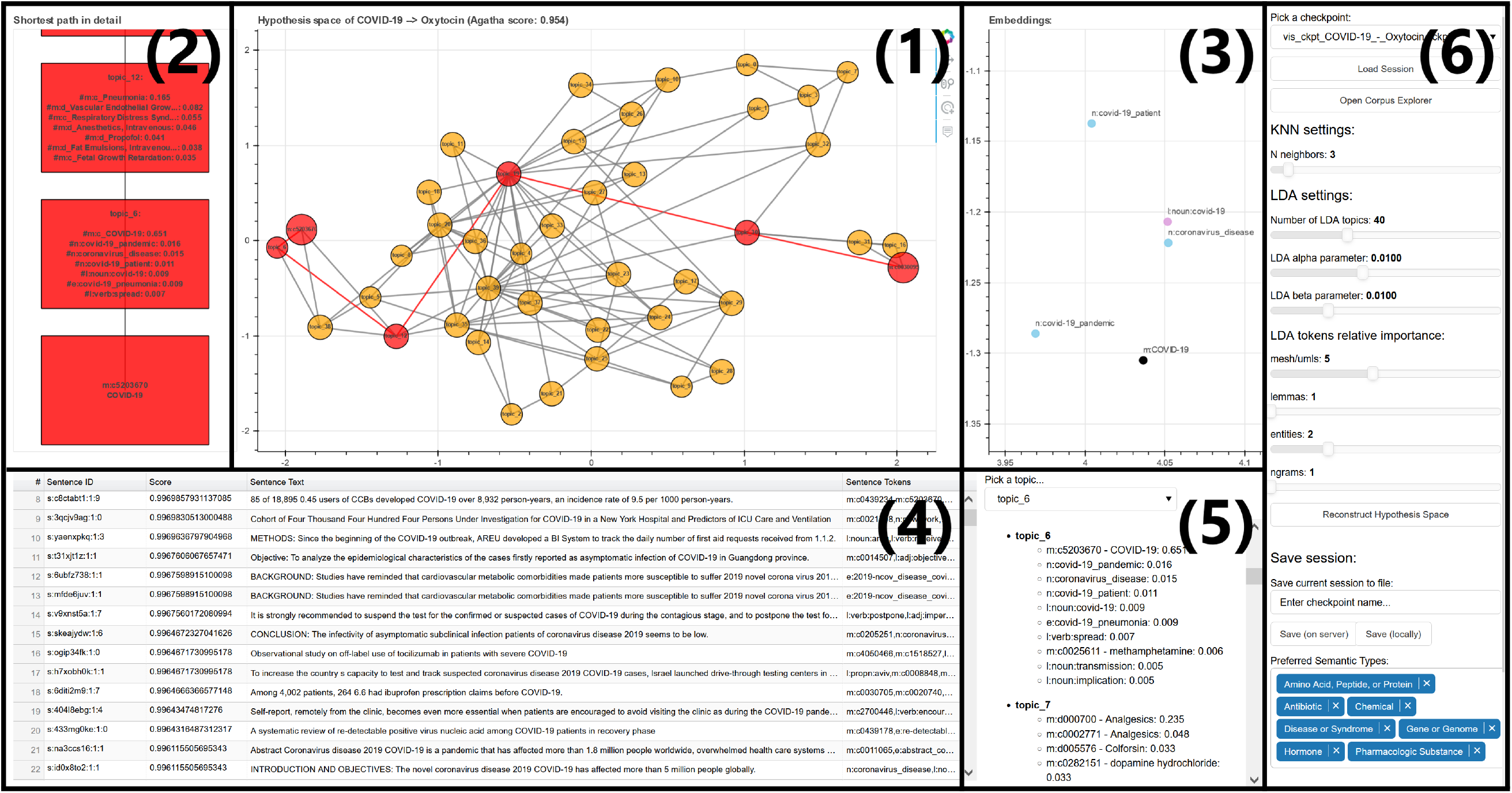
AGATHA Semantic Visualizer main window. Interface elements are numbered and described in Interface Walkthrough section. Image was rotated for better readability.

The first component is a computational server, which runs all computationally expensive tasks. This server contains an AGATHA instance and it is used to run visualization preprocessing stage and generate large-scale hypotheses ranking on demand. The example of the ranking containing more than 130.000 calculated pairwise scores related to COVID-19 queries is included in the system user guide shared with all domain experts reviewing the system.

The next part of the pipeline is a query submission form, which does not require a dedicated web server as part of the system and is hosted on Google servers for convenience.

The last component is a visualization web server, where the web interface is run. This web server contains a Bokeh [9] server instance, and all user sessions are independent from each other. It is used only for visualization/exploration purposes and for reconstructing the topical network on demand. All the functions that are available through the web-based GUI are performed on the visualization web server.

Pipeline is designed in this manner to isolate computationally intensive steps from the rest of the system. Generation visualizations occurs on demand meaning that powerful servers are not required to be utilized for the results displaying. Visualization server does not have significant system requirements, because the size of the data is significantly reduced during the preprocessing step.

### 4.2 Interface Walkthrough

#### Topical hypothesis space (1)

the “core” component of Agatha Hypothesis Space window. It shows how topics are arranged in latent semantic space and what are spatial relationships between source and target terms and topics related to them.

#### Shortest path in detail (2)

This pane contains a shortest path from the Topical hypothesis space pane, but in greater detail, such that a domain expert could see what each topic contains and how the transition between source and target terms occurs through the topics between them.

#### Embeddings pane (3)

displays the latent representation of terms within the topics participating in the highlighted shortest path, so there is a visual representation of not only how topics are arranged in the latent space, but also the terms, belonging to these topics. This window also illustrates how well graph embeddings represent concepts similarity.

#### Sentences pane (4)

shows a list of sentences associated with a specific topic and its confidence score. To pick a topic, one can use topics pane or click on a topic inside Topical Hypothesis Space.

#### Topics pane (5)

represents a list of all available topics generated from the corpus. A domain expert can scroll through it, pick a topic they are interested to construct a shortest path through and see what kind of chain of topics a system proposes, when an expert would like to see how the source term is connected to the target term through a specific midpoint.

This functionality is added to make use of *the entire topical space* and not just the shortest path suggested by the algorithm. While it is true that the default shortest path is the “best” way to connect source and target terms in terms of their spatial proximity in the embedding space, it sometimes may not contain enough contextual information about potentially relevant topics. A path through an intermediate topic is constructed by concatenating two shortest paths: first from the source term to the intermediate node and second from the intermediate node to the target term. If a selected topic is not relevant and has a significant distance from the default shortest path, a redundant loop may be present in the resulting path.

#### Settings pane (6)

allows a user to fine-tune the representation of learned topics and adjust the parameters of topic extraction process.

### 4.3 BioCreative VII IAT & User Feedback

This work is presented at BioCreative VII competition Interactive Text Mining Track (IAT), where it was evaluated by the experts involved in biomedical research and related fields. The track is motivated by formal evaluation of interactive tools and providing user feedback on text mining systems made for specific purposes. This year the main focus of IAT was COVID-19-related text mining tools.

We note that AGATHA is a general purpose hypothesis generation system not limited by any specific biomedical subdomain. It was recently enhanced with COVID-19 dataset CORD-19 [10], which made possible running queries involving COVID-19-associated terms and discovering novel connections, such as *COVID-19 - Oxytocin* [11]. Some of these connections were included in the default set of available visualizations.

The experts were offered to participate in guided and exploratory activities. The activities included working with different system functions, such as inspecting term-topical paths, reconstructing topical network, adjusting the parameters of KNN algorithm/LDA topic modeling, investigate the obtained document clustering and saving results. Upon activities completion, the experts were suggested to participate in a survey to formally evaluate the system.

The survey includes 32 questions, from them 26 questions are directly associated with the system. From these 26 questions 13 are rank-based and 13 are free-text/binary. We provide the boxplot of the responses to 12 rank questions having 1−5 scale in figure 4. The question *“How likely is it that you would recommend this system to a colleague performing COVID-19 related research?”* was on a scale of 1 − 10, where 1 = not at all likely and 10 = extremely likely. This question resulted in the mean value of 6, with the most frequent response of 7.

**Figure 4:**
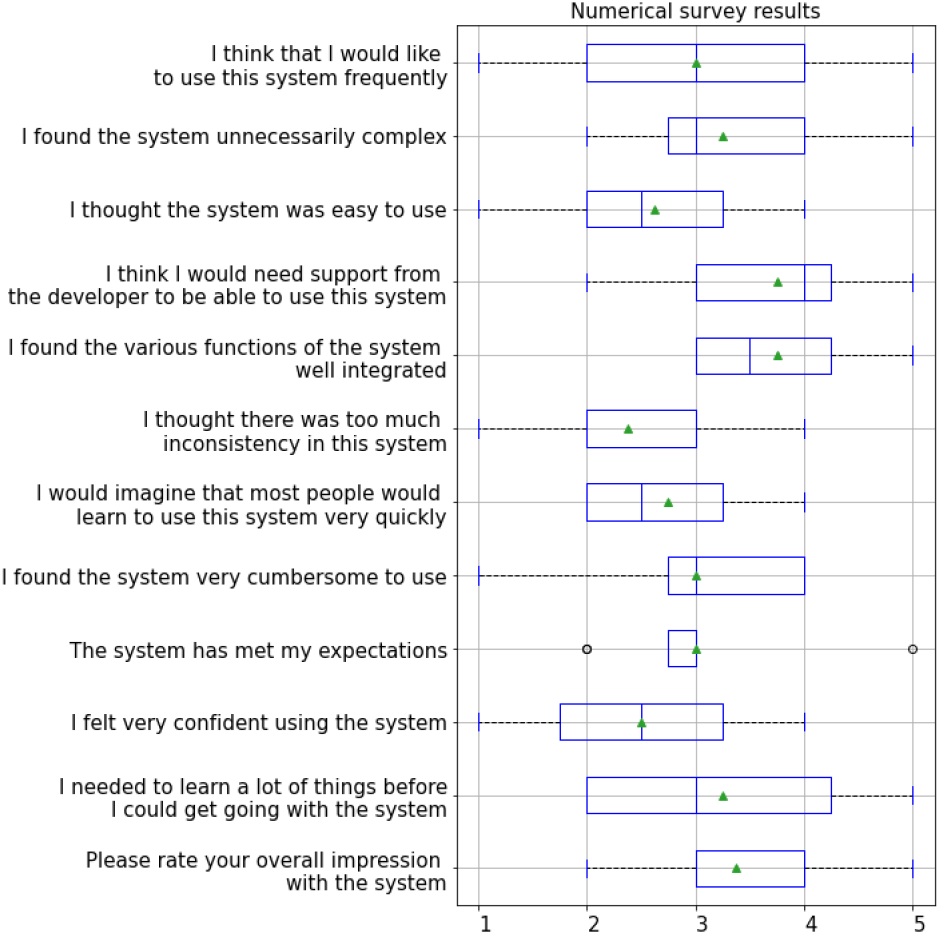
Boxplot of provided user feedback (rank-based questions, number of respondents: 8). Scale used: 1−5, where 1 was “strongly disagree” and 5 was “strongly agree”^2^.

Most respondents admit that they find the results produced by the system useful. Some respondents found results interpretation somewhat challenging due to specifics of topic modeling approach in general. Most of the comments and suggestions were addressed to the interface design and features, which some users found not intuitive enough. However, the overall approach to visual hypothesis exploration was generally well received.

Based on the feedback provided by the domain experts we plan to introduce a number of improvements. For example, interface changes (such as clickable PubMed links and built-in hints for parameter tuning) are expected to help domain experts to get better user experience. In addition, we observe that sentence-level granularity might not always be enough from the interpretability perspective, and more contextual information may be required. To add more clarity to the resulting topics (and, to the document clustering, respectively), it is possible to perform topic modeling on full abstracts (similar to the MOLIERE system) instead of individual sentences. This will not only introduce significantly more cohesive information for the visualizer input (individual sentences sometimes perform poorly for the topic modeling purposes), but will also make the final representation have less in common with the original semantic network, which is used for inference purposes.

## Footnotes

1 https://lhncbc.nlm.nih.gov/ii/tools/MetaMap/documentation/SemanticTypesAndGroups.html

2 With two exceptions: *How easy was it to format and input data into this tool?* 1 = not at all easy; 5 = extremely easy *Please rate your overall impression with the system*. 1 = very negative; 5 = very positive

## Notes

https://github.com/IlyaTyagin/AgathaSemanticVisualizer

### Competing Interest Statement

The authors have declared no competing interest.

